# Oxidative stress and loss of Fe-S proteins in Friedreich ataxia induced pluripotent stem cell-derived PSNs can be reversed by restoring FXN expression with a benzamide HDAC inhibitor

**DOI:** 10.1101/221242

**Authors:** Amelié Hu, Myriam Rai, Simona Donatello, Massimo Pandolfo

**Author notes:** To whom correspondence should be addressed at: Massimo Pandolfo - Laboratory of Experimental Neurology, Université Libre de Bruxelles CP-601, Route de Lennik 808, 1070 Brussels, Belgium.

## Abstract

Epigenetic suppression of frataxin (FXN) expression caused by the presence of expanded GAA repeats at the *FXN* locus is the key pathogenic event in Friedreich ataxia (FRDA), a recessive neurodegenerative and systemic disease. FXN is involved in iron-sulfur (Fe-S) cluster biogenesis in mitochondria, its deficiency causes multiple Fe-S protein deficiencies, mitochondrial dysfunction and oxidative stress. Primary sensory neurons (PSNs) in the dorsal root ganglia (DRGs) are the most vulnerable cells in FRDA, whose abnormal development and degeneration leads to the onset and early progression of ataxia. We generated PSNs from induced pluripotent stem cells (iPSCs) from FRDA patients and showed that they recapitulate the key pathogenic events in FRDA, including low FXN levels, loss of Fe-S proteins and impaired antioxidant responses. We also showed that FXN deficiency in these cells may be partially corrected by a pimelic benzamide histone deacetylase inhibitor, a class of potential therapeutics for FRDA. We generated and validated a cellular model of the most vulnerable neurons in FRDA, which can be used for further studies on pathogenesis and treatment approaches.

## Introduction

Friedreich's ataxia (FRDA) is an autosomal recessive multisystem disorder with prominent neurological manifestations, often associated with hypertrophic cardiomyopathy, skeletal abnormalities and carbohydrate intolerance (Cnop et al., 2013; Pandolfo and Manto, 2013; Parkinson et al., 2013). In almost all cases, the underlying genetic mutation is the hyperexpansion of a GAA repeat in the first intron of the *FXN* gene, encoding frataxin (FXN). The expanded repeats suppress *FXN* expression by triggering chromatin condensation. Accordingly, repressive epigenetic marks are found in the gene, such as loss of acetylation of histones H3 and H4 and trimethylation of histone H3 lysine 9 (H3K9m3)(Herman et al., 2006; Soragni et al., 2014). A specific class of histone deacetylase (HDAC) inhibitors (HDACi) can reverse those changes and restore *FXN* expression in cells from FRDA patients and animal models, providing a potential treatment for the disease(Soragni et al., 2014). Target specificity of benzamide HDACi for the class I HDACs HDAC1/2 and HDAC3, and slow-on slow-off kinetics appear to be critical for FXN upregulation (Chou et al., 2008; Xu et al., 2009). Importantly, in FXN upregulation is obtained at doses that cause minimal overall gene expression changes (Coppola et al., 2011), suggesting that toxicity due to gene expression dysregulation is unlikely. However, restoring FXN levels can be therapeutic only as long as pathogenic changes induced by FXN deficiency are reversible. FXN is a component of the protein complex that synthesizes iron-sulfur (Fe-S) clusters in the mitochondrial matrix. Its deficiency leads to multiple metabolic abnormalities due to loss of Fe-S proteins activities, altered iron homeostasis and oxidative stress. We recently showed that these abnormalities are found in neurons differentiated from induced pluripotent stem cells (iPSC) from FRDA patients and are reversible when FXN levels are normalized by treatment with a benzamide HDAC inhibitor(Codazzi et al., 2016). In that study we obtained and characterized neurons with a cortical phenotype, but primary sensory neurons (PSN) in the dorsal root ganglia (DRGs) are the most vulnerable and earliest affected cells in FRDA. Here we report the generation of PSNs from FRDA patients iPSCs, the initial characterization of their biochemical and molecular phenotype and the restoration of FXN levels by benzamide HDACi.

## Materials and Methods

### Neurosphere culture and differentiation

Experiments were carried out using induced pluripotent stem cells (iPSC) obtained by reprogramming skin fibroblasts from two FRDA patients and two healthy controls. The generation and validation of these iPSC lines, and their differentiation into neurons and cardiomyocytes were previously described(Hick et al., 2013). Briefly, iPSC colonies were treated with the bone morphogenetic protein (BMP) inhibitor noggin for 14 days, generating neurospheres that were then mechanically transferred and cultured in NBN media NBN media (Neurobasal media A+B27+N2) with FGF-2 and EGF.

### Culture of iPSC-derived primary sensory neurons

Experiments were carried out using induced pluripotent stem cells (iPSC) obtained by reprogramming skin fibroblasts from two FRDA patients (FA 1 and FA 2) and one healthy control (CT 1). We previously reported the generation and validation of these iPSC lines (Hick et al., 2013). Neural induction was performed as previously described(Chambers et al., 2012) with few modifications. Briefly, iPSC colonies were treated with 100μM LDN-193189 (Sigma; USA) and 10μM SB431542 (Tocris bioscience; UK) in iPSC medium (Knockout DMEM, 20% Knockout Serum Replacement, 1mM L-glutamine, 100μM MEM non-essential amino acids, Penicillin-Streptomycin and 0.1 μM ß-mercaptoethanol), on days 0 through day 5. Cells were fed every day and on day 4, N2 media was added in increasing 25% increments every other day (100% N2 on day 10). Primary sensory induction was started with the addition of 3μM CHIR99021, 10μM SU5402 and 10μM DAPT (Tocris bioscience; UK) on days 2 through day 10. On day 11, cells were mechanically transferred and cultured in NBN medium (Neurobasal-A medium, 2% B27 supplement, 1% N2 supplement), in the presence of Fibroblast Growth Factor 2 (FGF-2) and Epidermal Growth Factor (EGF) (both 20μM/ml; Peprotech, UK), which allowed the formation of neurospheres. Neural precursors were maintained and propagated as neurospheres by the cutting method and by replacing 50% of the medium every 2 days. For neuronal differentiation, neurospheres were plated on laminin-coated glass coverslips in NBN media in presence of 25 ng/ml Brain-Derived Neurotrophic Factor (BDNF), Glial cell line-Derived Neurotrophic Factor (GDNF) and Nerve Growth Factor (NGF). Cells were fed every 2-3 days and differentiated during 2 weeks.

### Cell treatments

Differentiated cells were treated with DMSO or 5μM of N-(6-(2-aminophenylamino)-6-oxohexyl)-4-methylbenzamide pimelic diphenylamide (HDACi 109) during 24 hours and 72 hours(Soragni:2014hk; Codazzi et al., 2016). Cells were rinsed in PBS on ice, and collected as dry-pellet for proteins extraction or lysed in RLT-ß-mercaptoethanol solution for RNA extraction.

### Immunofluorescence

Differentiated cells cultured on glass coverslips coated with 0.4% laminin, were rinsed with PBS and fixed in PBS containing 4% paraformaldehyde for 15 minutes at room temperature. Cells were permeabilized with 0.1% Triton X-100 for 15 minutes and non-specifics sites were blocked with Blocking Buffer (PBS, 5% NGS, 5% NHS) for 30 minutes at room temperature. They were then incubated overnight at 4°C with primary antibodies diluted in blocking buffer. Primary antibodies, rabbit anti-Brn3a (Millipore; AB5945) and rabbit anti peripherin (Abcam, ab99942), diluted 1:500 and 1:1000 respectively, were used as PSNs markers. The secondary antibody anti-rabbit (1:1000; Invitrogen) was incubated at room temperature for 1 hour. Differentiated cells were incubated 5 minutes with 10μM/ml of DAPI (4-,6-diamidino-2-phenylindole) at room temperature, to counterstain cell nuclei. Then, cells were mounted on microscope slides using Fluorsave (Calbiochem, Germany). Results were visualized by AxioImager Z1 microscope (Zeiss, Jena, Germany), and exported as TIFF files to be analyzed with ImageJ program.

### Quantitative Real time PCR

Total RNA was isolated from differentiated cells by using of RNeasy Mini Kit (Qiagen). RNA samples were treated with RNase-Free DNase Set (Qiagen) and quantified afterwards by measuring the optical density (NanoDrop ND-1000 Spectrophotometer, NanoDrop Technologies). We performed one-step qRT-PCR using a 7500 Fast Real-Time PCR system with MultiScribe Reverse Transcriptase and Power SYBR Green (Applied Biosystems). Primers used for *FXN* were 5’-CAGAGGAAACGCTGGACTCT-3’ and 5’-AGCCAGATTTGCTTGTTTGG-3’, for *SOD2* were 5’- TTTCAATAAGGAACGGGGACAC-3’ and 5’-GTGCTCCCACACATCAATCC-3’ and for *NRF2* were 5’-TTCCCGGTCACATCGAGAG-3’ and 5’- TCCTGTTGCATACCGTCTAAATC-3’. Cells RNA was standardized by quantification of GAPDH. Data are normalized to the *FXN* or *SOD2* or *NRF2* mRNA level in CT cells (100%).

### Western blotting

Total extracts were obtained by homogenizing cells in 20mM Tris-HCl, pH 7.5, 150 mM NaCl, 1mM Na2EDTA, 1mM EGTA, 1% NP-40, 1% Sodium deoxylcholate, 2.5mM sodium pyrophosphate, 1mM β-glycerophosphate, 1mM Na_3_VO_4_, 1μg/ml leupeptin and 1mM PMSF. Protein concentration was determined using the Pierce BCA protein assay kit. Proteins (10 μg) were denatured for 5 minutes at 100°C in 2 × loading buffer and resolved by electrophoresis on a 12% polyacrylamide SDS-PAGE gel. Membranes were blocked for 1 hour at room temperature in Odyssey blocker. Membranes were incubated overnight at 4°C in antibodies anti-FXN (1:1000; Sc25820, Santa Cruz), anti-aconitase (1:500; ab71440, Abcam), anti-NDUFS3 (1:1000; PA5-29747, Thermo Scientific), anti-Lipoic Acid (1:1000; 437695, Calbiochem), anti-ISCU (1:1000; Sc28860, Santa Cruz), anti-IRP1 (1:1000; Sc14216, Santa Cruz), anti-Citrate synthase (1:1000; Sc390693, Santa Cruz), anti-SOD2(1:1000; Sc30080, Santa Cruz), anti-Nrf2 (1:250; ab62352, Abcam) anti-GAPDH (1:1000; cb1001, Calbiochem) and anti-α-Tubulin (1:10000; 070M4755, Sigma). The membranes were incubated for 1 hour in secondary antibody (Goat anti-mouse, 1:15000, 926-32220, LI-COR; Donkey anti-goat, 1:10000, 925-68074, LI-COR; Donkey anti Rabbit, 1:15000, 926-32213, LI-COR) at room temperature and scan using LI-COR Odyssey western blotting detection system. Band intensities were quantified by using the ImageJ software. GAPDH, α-Tubulin and CS (for some experiments involving mitochondrial proteins) were used for normalization. Each protein was quantified in four biological replicates.

### Statistical analysis

Data are presented as mean ± SD. Comparisons between CT and FA 1 and FA 2 were made using non-parametric test, Kruskal-Wallis, followed by Dunn post-hoc test using the Prism Software (GraphPad Software, San Diego, CA, USA). *p*-values less than 0.05 were considered significant.

## Results

### Generation of PSNs derived from iPSC

In order to obtain faithful cell models that recapitulate the pathophysiological features of FRDA, we generated PSNs from iPSCs from two controls (CT) and two FRDA patients (FA 1 and FA 2), as previously described(Chambers et al., 2012) with few modifications (see Materials and Methods). To induce neuronal differentiation, iPSCs were treated on day 1 to 5 with two inhibitors of SMAD pathway, bone morphogenetic protein (BMP) inhibitor LDN-193189 and SB431542. On day 2 to 10, three small molecules were added: SU5402, a potent inhibitor of vascular epithelial growth factor (VEGF), fibroblast growth factor (FGF) and platelet-derived growth factor (PDGF) tyrosine kinase signaling(Sun et al., 1999); CHIR99021 which can act as a WNT agonist by selectively inhibiting glycogen synthase kinase-3β (GSK-3β) and thereby stabilizing β-catenin(Bennett et al., 2002); and DAPT which is a γ-secretase inhibitor that blocks Notch signaling(Dovey et al., 2001). On day 11, neural precursors were mechanically isolated and cultured in suspension as neurospheres. After 2 weeks in culture of differentiation conditions, FRDA and CT cells were positive for the PSNs markers Brn3a (Brain-specific homeobox/POU domain protein 3A) and peripherin (type III Intermediate filament) (Figure 1A). In these cells, levels of Brn3a, peripherin and islet1 (Insulin gene enhancer protein ISL-1) were much higher than levels in cortical neurons generated from the same iPSCs as previously described(Hick et al., 2013), with a 5-fold enrichment for Brn3a and Peripherin, and a 4-fold enrichment for Islet1 (Figure 1B).

**Figure 1.**
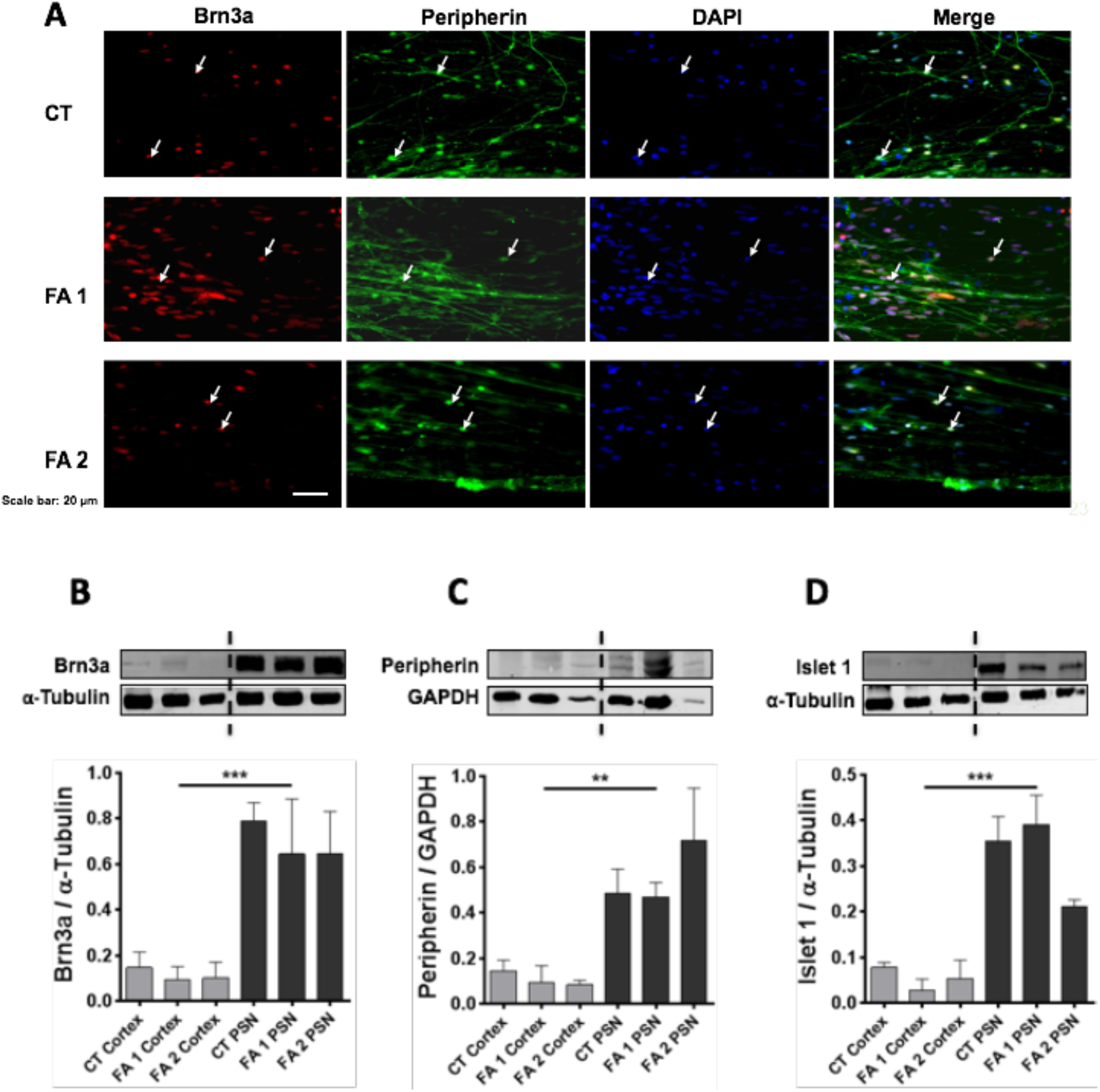
Characterization of FRDA and control iPSC-derived PSNs. Brn3a and Peripherin expression were assessed by immunocytochemistry (A) and Brn3a, Peripherin and Islet1 by western blot (B-D) in iPSC-derived primary sensory neurons from two FRDA patients (FA1 and FA) and one healthy control (CT1). (A) Immunocytochemistry for Brn3a and Peripherin. (B-D) Western blot analysis (n=3 independent experiments). *FRDA versus CT, **p<0.01, ***p<0.005, by the Kruskal-Wallis test, a nonparametric analogue to ANOVA, followed by Dunn's *post hoc* test. P-values less than 0.05 were considered significant.

### FXN and Fe-S proteins expression is decreased in FRDA primary sensory neurons

PSNs from FRDA patients had significantly reduced FXN protein (Figure 2A) and mRNA levels compared to CT cells (Figure 2B)Levels of ISCU, a component of the multiprotein complex that assembles Fe/S clusters in the mitochondrial matrix, along with NFS1, ISD11 and FXN(Schmucker et al., 2011), were also reduced in FRDA primary sensory neurons, though not to the same extent as FXN (Figure 2C). Levels of Fe-S proteins (Aconitase and NDUFS3) were 60-70 % lower in FRDA PSNs compared to CT (Figure 2D), while expression of citrate synthase, a mitochondrial enzyme not containing an Fe/S cluster and commonly used as marker for mitochondrial matrix proteins, was unchanged (Figure 2E).

**Figure 2.**
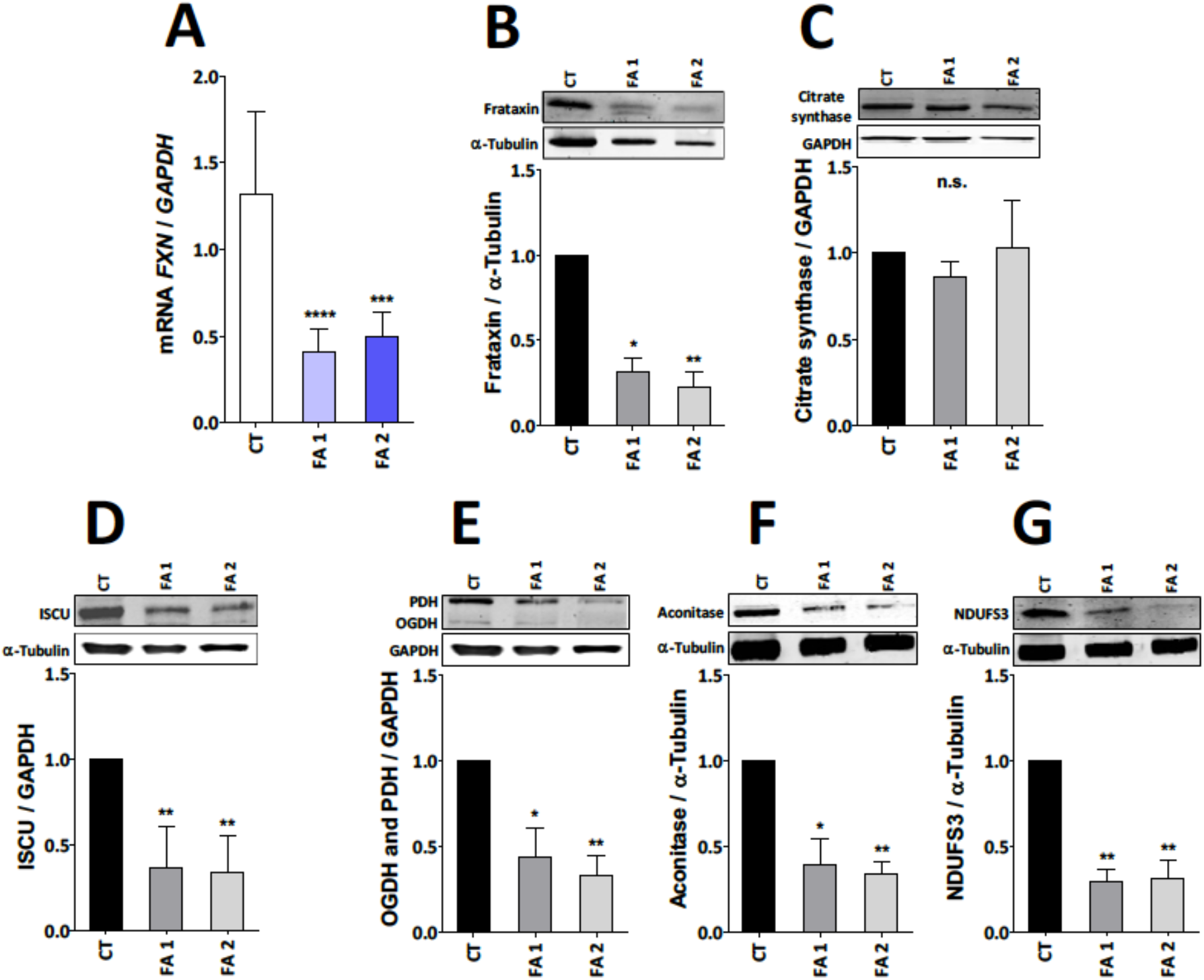
FXN mRNA and protein levels and Fe-S proteins expression decreased in FRDA PSNs. FXN mRNA levels were determined by quantitative real-time RT-PCR (A), FXN and Fe-S proteins expression were determined by western blot (B-G), in iPSC-derived PSNs from two FRDA patients (FA1 and FA2) and one healthy control (CT1). (A) FXN mRNA levels. (B-E) Western blot analysis (n=6 independent experiments). *FRDA versus CT, *p<0.05, **p<0.01, ***p<0.005, ****p<0.001, by the Kruskal-Wallis test, a nonparametric analogue to ANOVA, followed by Dunn's *post hoc* test.

Aconitase is a mitochondrial [4Fe-4S] enzyme implicate in a Krebs cycle and NDUFS3 (NADH dehydrogenase iron sulfur protein 3) is a subunit of complex 1. Furthermore, levels of lipoic acid-bound PDH (pyruvate dehydrogenase) and OGDH (2-oxoglutarate dehydrogenase) were similarly reduced. Lipoic acid is synthetized by the [4Fe-4S] enzyme lipoic acid synthase(Navarro-Sastre et al., 2011), so its abundance is reduced when Fe-S cluster biogenesis is impaired.

### FXN deficiency alters oxidative stress responses in FRDA primary sensory neurons

FRDA PSNs showed similar abnormal markers of increased oxidative stress as iPSC-derived FRDA cortical neurons(Codazzi et al., 2016). As observed in cortical neurons(Codazzi et al., 2016), mitochondrial superoxide dismutase 2 (SOD2) protein was 2-3 fold higher in FRDA than in CT neurons (Figure 3F), but its activity was impaired and its mRNA was decreased by 50% (Figure 3B). Levels of the master regulator of antioxidant responses nuclear factor (erythroid-derived 2)-like 2 (NRF2) mRNA and protein were also decreased, (Figure 3C and 3E).

**Figure 3.**
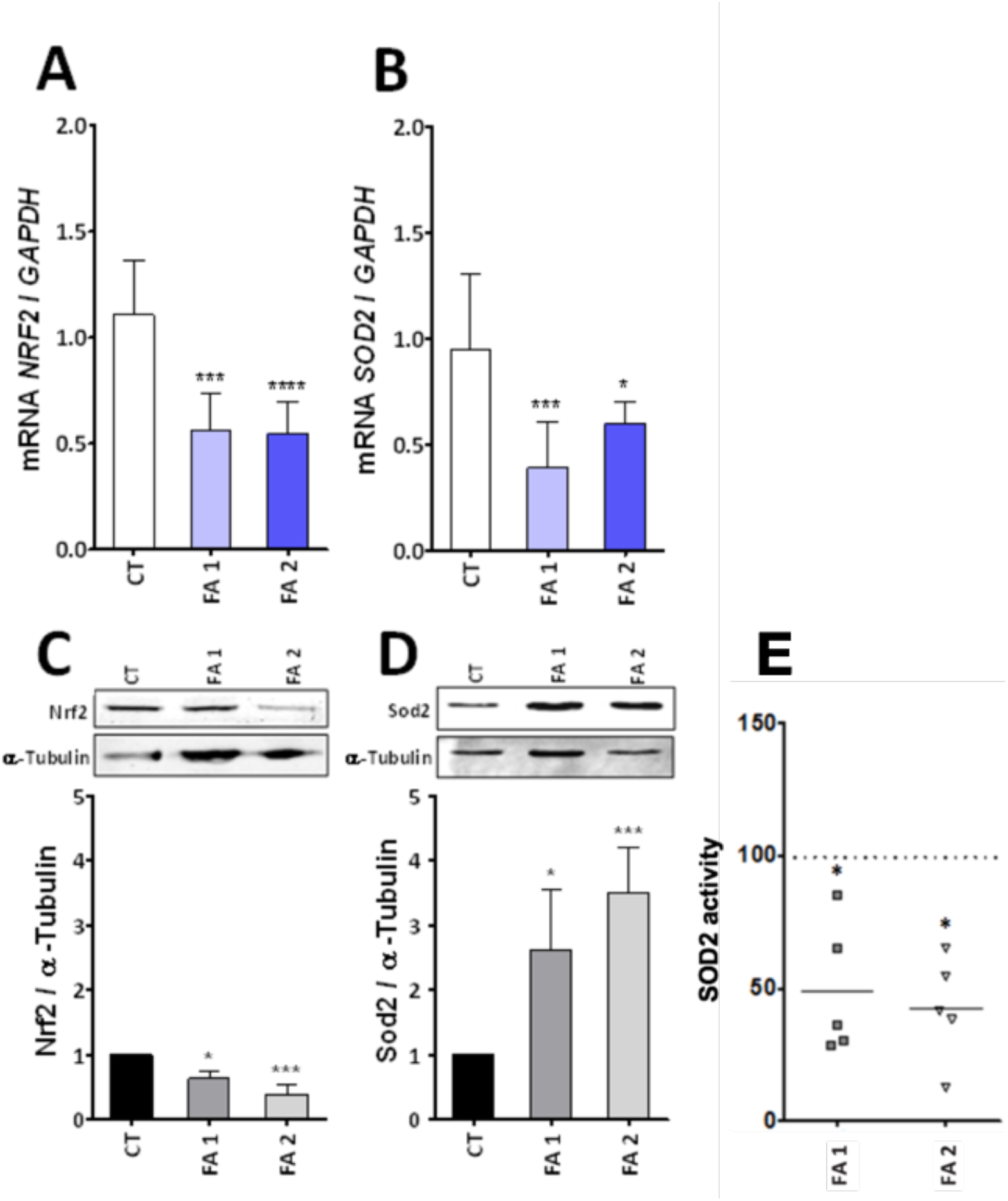
Mitochondrial stress induced by frataxin deficiency in primary sensory neurons FRDA. NRF2 and SOD2 mRNA levels by quantitative real-time RT-PCR (A and B), NRF2 and SOD2 protein levels by western blot analysis (C and D), SOD2 activity expressed as percentage of CT levels (E) in iPSC-derived PSNs from two FRDA patients (FA1 and FA2) and one healthy control (CT1). (A and B) NRF2 and SOD2 mRNA levels analysis. (C and D) Western blots analysis (n=6 independant experiments). *FRDA versus CT, *p<0.05, **p<0.01, ***p<0.005, ****p<0.001, by the Kruskal-Wallis test, a nonparametric analogue to ANOVA, followed by Dunn's *post hoc* test.

### Effect of benzamide HDACi on FXN mRNA and protein

Treatment of FRDA PSN with 5μM of the benzamide HDAC inhibitor 109 for 24 h or 72 h resulted in significant increase of frataxin mRNA and protein (Figure 4C and 4D), while there was no effect in CT cells (Figure 4A and 4B).

**Figure 4.**
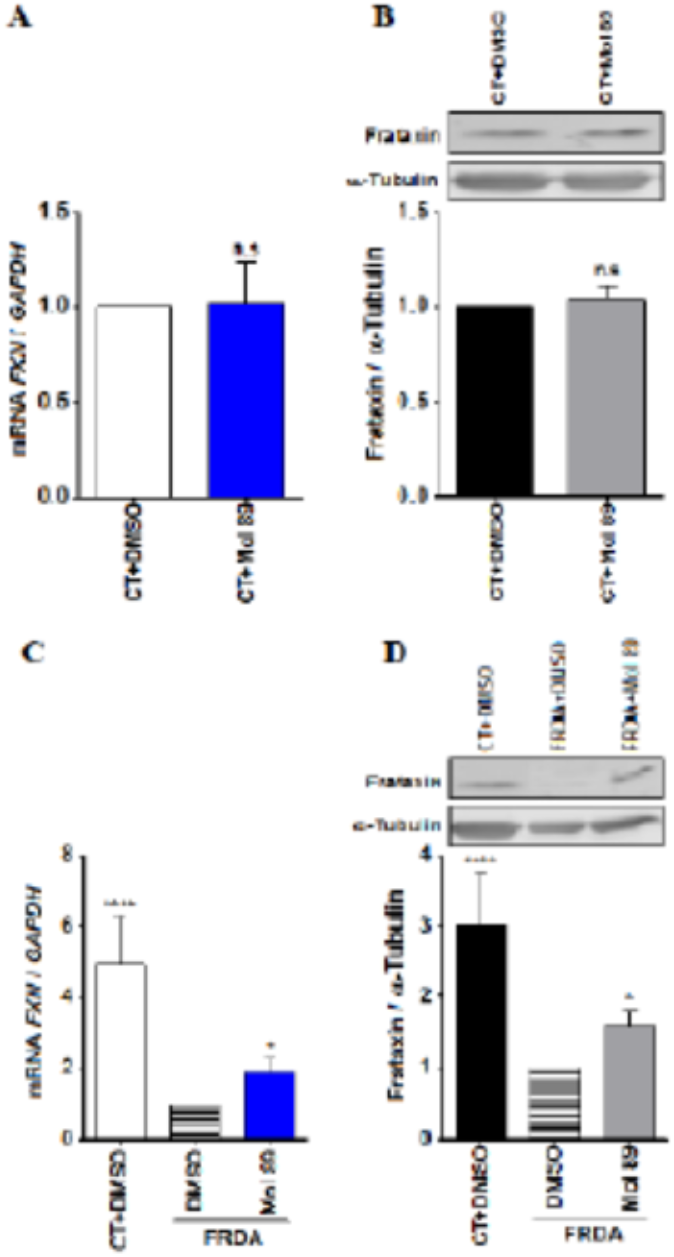
Effect of HDACi 109 in FRDA PSNs. Treatment of iPSC-derived PSNs from two FRDA patients (FA1 and FA2) and one healthy control (CT1) with 5μM of the benzamide HDAC inhibitor RG2033 for 24 h or 72 h had no effect in CT cells (A and B), but resulted in significant increase of FXN mRNA (C) and protein (D) by the Kruskal-Wallis test, a nonparametric analogue to ANOVA, followed by Dunn's *post hoc* test. *FRDA versus CT, *p<0.05, **p<0.01, ***p<0.005.

## Discussion

At symptom onset and during early progression, FRDA is essentially a sensory neuronopathy affecting large PSNs in dorsal root ganglia (DRG), causing afferent ataxia, proprioceptive loss, and loss of tendon reflexes. Later, cerebellar ataxia, dysarthria, oculomotor abnormalities, pyramidal weakness (Koeppen and Mazurkiewicz, 2013; Pandolfo and Manto, 2013), and auditory and visual loss (Fortuna et al., 2009; Rance et al., 2012) characterize disease progression. Neuropathological studies of DRGs, dorsal roots and spinal cord of FRDA patients even suggest that PSNs may already be affected during development(Koeppen et al., 2017a; Koeppen et al., 2017b), and continue to degenerate in symptomatic individuals(Koeppen et al., 2016). We have no explanation for such an exquisite sensitivity of large PSNs to FXN deficiency. The possibility to generate PSNs from iPSCs, as reported here, provides an opportunity to study how FXN deficiency affects these cells during and after their differentiation. We could show that these cells present the basic biochemical and molecular changes expected to be caused by FXN deficiency, including deficits in Fe-S proteins and abnormal antioxidant responses. In particular, we confirmed dysregulation of SOD2, which shows low activity and mRNA levels, evidence of blunted antioxidant responses(Chantrel-Groussard et al., 2001; Jiralerspong et al., 2001). SOD2 protein levels were instead increased, as we had observed in iPSC-derived FRDA cortical neurons(Codazzi et al., 2016), suggesting that the protein may be dysfunctional, as previously reported in a yeast model and attributed to Fe toxicity(Irazusta et al., 2010). We also found low mRNA and protein levels of NRF2, a master regulator of antioxidant responses known to be impaired in other FRDA models(Paupe et al., 2009). This finding suggests that NRF2 activators, now being tested in a clinical trial as potential treatment for FRDA, may also target PSNs in DRGs.

Finally, we demonstrated the efficacy of benzamide HDACi in upregulating FXN in FRDA PSNs, further supporting the clinical development of this category of drugs as FRDA therapeutics.

The validation of this model provides a resource for further studies on the specific vulnerability of FRDA PSNs by investigating differences between control and FRDA cells during and after full differentiation in terms of morphology, subpopulation specific markers, gene expression profiles, and electrophysiological properties, as well as to test novel therapeutic approaches for FRDA.

## Acknowledgements

Financial support was from the Friedreich Ataxia Research Alliance (FARA) to MP; the European Commission grant HEALTH-F2-2010-242193 (European Friedreich's Ataxia Consortium for Translational Studies - EFACTS) to MP; the National Ataxia Foundation (NAF) research grant 2014 to SD.

